# Dissecting genomic regions and candidate genes for pod borer resistance and component traits in pigeonpea minicore collection

**DOI:** 10.1101/2025.08.04.668421

**Authors:** Abhinav Moghiya, R.S. Munghate, Vinay Sharma, Suraj Prashad Mishra, Jagdish Jaba, Shailendra Singh Gaurav, Sunil S. Gangurde, Namita Dube, Sagar Rangari, Rajib Roychowdhury, Prakash Gangashetty, Hari Chand Sharma, Manish K Pandey

## Abstract

**Background:** Pigeonpea is an important leguminous food crop primarily grown in tropical and subtropical regions of the world and is a rich source of high-quality protein. The biotic (weed, disease, and insect-pests) and abiotic stresses have significantly reduced the production and productivity of pigeonpea. *Helicoverpa armigera*, also known as pod borer, is a major pest in pigeonpea. A substantial investigations needed for comprehending the genetic and genomic underpinnings of resistance to *H. armigera*. Genetic improvement by genomics-assisted breeding (GAB) is an effective approach for developing high yielding *H. armigera* resistance cultivars. Still yet, no genetic markers and genes linked to this key trait have been detected in pigeonpea. In this context, a set of 146 pigeonpea minicore accessions were evaluated for four *H. armigera* resistance component traits namely pod borer resistance (PBR), days to 50% flowering (DF), days to maturity (DM), and grain yield (GY) for three consecutive seasons under field conditions.

**Results:** Phenotypic data of pod borer resistance and component traits, along with the whole-genome resequencing (WGRS) data for 4,99,980 SNPs, were utilized to perform multi-locus genome-wide association study (GWAS) analysis. Two models (SUPER, and FarmCPU) detected 14 significant marker-trait associations (MTAs) for PBR and three component traits. The MTAs with significant effect were mainly identified on chromosomes CcLG02, CcLG04, CcLG05, CcLG07 and CcLG11. These MTAs subsequently delineated with key candidate genes associated with pod borer resistance (*Probable carboxylesterase 15, microtubule-associated protein 5, FAR1-RELATED SEQUENCE*, and *Omega-hydroxypalmitate O-feruloyl transferase 4*), days to maturity (*RING-H2 finger protein ATL7* and *Leucine-rich repeat receptor-like protein kinase*), and grain yield (*Secretory carrier-associated membrane protein*, and *Glutaredoxin-C5 chloroplastic*).

**Conclusion:** These research findings reported significant MTAs and candidate genes associated with pod borer resistance and component traits. Further lab-based pod bioassay screening identified four minicore accessions, namely ICP 10503, ICP 655, ICP 9691 and ICP 9655 (moderately resistance genotypes), showing least damage rating and larval weight gain %, compared to the susceptible checks. After validating the significant MTAs, the associated SNP markers can be effectively utilised in indirect selection, which offers potential gains for such quantitative traits with low heritability and can improve insect management more sustainably. The significant MTAs, candidate genes and resistant accessions, reported in this study may be utilized for the development of pod borer-resistant pigeonpea varieties.

## Introduction

Pigeonpea [*Cajanus cajan* (L.) Millsp.] is an important food legume crop in arid and semiarid regions of Asia and Africa. It is grown on 5.7 million hectares worldwide, with a production of 4.9 million tons (FAO, 2024). India, along with Malawi, Tanzania, Kenya, Uganda and Myanmar, is a leading producer contributing 78% of the global pigeonpea production. As one of the five major edible legumes, pigeonpea is used for edible purpose, animal feed, and firewood. It is an important source of protein, often used to supplement cereal-based diets (Kinhoégbè et al., 2022). Climate change presents a substantial risk to worldwide pigeonpea production, impacting both its nutritional quality and its ability to withstand various abiotic and biotic stresses. In India, pulses are vulnerable to approximately 150 insect pest species (Seetharamu et al., 2020), and globally, around 38 species of Lepidopteran insect harm pigeonpea (Shanower et al., 1999). Among the most damaging biotic stresses is the pod borer, *Helicoverpa armigera*, which severely affects crop growth and yield (Ghosh et al., 2017). Although pesticides can control pod borer complex (PBC), excessive use of chemical insecticides has resulted in insect susceptibility, secondary pest outbreaks, detrimental impacts on biodiversity and negative environmental effects (Ambidi et al., 2021; Jaba et al., 2023). Therefore, developing pigeonpea varieties that are resistant to *H. armigera* is seen as the most effective solution to reduce pesticide use. Despite extensive screening of various pigeonpea genetic resources across Asia and Africa, no strong resistance against pod borer has been reported (Kambrekar 2016). However, partial resistance has been reported in some cultivated genotypes, which has been utilized in pigeonpea breeding programs. While wild pigeonpea species confer higher resistance to pod borer (PBR) compared to cultivated sources, transferring these resistance genes to cultivated varieties is limited to only a few wild species due to cross-incompatibility (Sharma, 2016; Singh et al., 2020). In earlier investigations, the minicore collection was screened, showing moderate resistance levels to the pod borer (Sharma et al. 2025 Unpublished). This data has now been utilised for conducting GWAS and facilitating gene discovery.

The development and use of genomic tools can facilitate selection of genotypes/breeding lines that are resistant to *H. armigera* using marker-assisted selection (MAS). However, there seems to be a lack of efforts in identifying candidate genes and markers. Molecular markers are important for facilitating the transfer of insect resistance genes into elite backgrounds, elucidating gene action, and minimising the negative effects of integrating undesirable genes from wild relatives due to linkage drag. Molecular breeding holds the potential to pyramid various sources of resistance that might not be efficiently selected by conventional breeding strategies due to phenotypic similarities, hence can increase resistance levels and potentially developing resistant varieties (Sharma & Crouch, 2004). Consequently, an innovative approach might be needed to integrate traditional approaches with MAS to develop systems preferable than either approach alone. Recent breakthroughs in pigeonpea genomics research have resulted in the development of draft and telomere-to-telomere reference genomes (Varshney et al., 2012, Garg et al., 2022, Liu et al 2024). Additionally, the accessibility of whole-genome sequencing (WGS) data (Varshney et al., 2017) and high-density Axiom *Cajanus* SNP arrays with 56K SNPs (Saxena et al., 2017) has significantly advanced genetic diversity, QTLseq and genome-wide association analysis studies. Genome-wide association study (GWAS) or association mapping has emerged as a important tool for identifying MTAs, candidate genes and associated markers (Gudi et al., 2024; Sharma et al., 2024). WGRS-based GWAS is effective for identifying associated genomic regions and candidate genes related with specific traits in various legume species, including pigeonpea (Varshney et al., 2012, Xu et al., 2017; Kang et al., 2019). Recent studies, detected MTAs for flowering time (Kumar et al. 2022); and antioxidant properties (Megha et al., 2024). Similarly, MQTLs were identified for agronomic trait, fertility restoration, disease resistance and seed quality traits (Halladakeri et al., 2023). This investigation utilized a multi-season phenotyping data generated on diverse minicore accessions to identify significant MTAs and candidate genes linked with pod borer resistance. We highlighted the importance of using various resistance sources against pod borer damage, emphasizing the relationships between component traits (phenology and grain yield) and resistance levels. These findings facilitate the development of pigeonpea varieties exhibiting improved resistance to pod borer.

## 1. Materials and methods

### 2.1 Plant material

This investigation used 146 accessions from the ICRISAT minicore collection (Upadhyaya et al., 2006), along with 2 checks (resistant check ICPL 332WR and susceptible check ICPL 87). Seed material were procured from the ICRISAT Genebank (https://genebank.icrisat.org/IND/Passport?Crop=Pigeonpea&Location=Passport&mc=Yes).

### 2.2 Field experiment, phenotyping for pod borer resistance and component traits

Phenotypic screening of 146 accessions, including the susceptible check ICPL 87 and the resistant check ICPL 332WR, was performed using a randomized block design with three replicates during Rainy 2007 (S1), Rainy 2008 (S2) and Rainy 2009 (S3) at ICRISAT-Patancheru, Hyderabad. Each plot consisted of 4 rows with a row spacing of 30 cm and a plant spacing of 10 cm within each row. Plots were separated by 1m alley. Five randomly selected plants from each genotype and replication were tagged for recording observation on pod borer under natural infestation at maturity stage. Pod borer resistance (PBR) was evaluated via a visual damage score on a scale of 1-9, where 1 indicates almost no damage (resistant), and 9 represents severe damage (highly susceptible) (**Supplementary Figure 1**), during the pod stage (Sujana et al., 2008). This was assessed alongside component traits such as, days to 50% flowering (DF), days to maturity (DM), and grain yield (GY) DF and DM were recorded on a per plant basis. The following pigeonpea descriptors (IBPGR and ICRISAT, 1993) were used to record the GY (g) per plant on five randomly selected representative plants per plot. The best four minicore accessions were subjected to a pod bioassay with artificial 3^rd^ instar larvae (Ambidi et al., 2021)

### 2.3 Statistical analyses of phenotypic data

The statistical analysis of the phenotypic data was performed via R Studio version 4.3.1 (http://www.rstudio.com/). The ‘FactoMineR’ package in R was used to perform Pearson correlation on replicated data from three seasons (Lê et al., 2008). The R package “phenotype” was used to calculate the BLUPs (Piepho et al., 2008).

### 2.4 DNA extraction and whole-genome resequencing (WGRS)

Genomic DNA was isolated from young leaves using the NucleoSpin® 96 Plant II (Macherey-Nagel) Kit. The quality was assessed through 0.8% agarose gel electrophoresis, and the amount was quantified via a Qubit® 2.0 fluorometer (Thermo Fisher Scientific Inc., USA) (Pandey et al., 2021). We generated libraries with a 500 bp insert size for all samples for WGRS, as detailed in Varshney et al. 2017. The fragments with insert sizes of around 500 bp were removed following separation on an agarose gel and then amplified by PCR. Furthermore, each library was subjected to sequencing on the Illumina HiSeq 2500 to generate paired-end reads. The raw reads were subjected to quality check using FastQC v0.11.8, and Trimmomatic v0.39 and poor reads (Phred score < 30, read length < 35 bp) and adaptor exhibiting contamination were eliminated, resulting in high-quality reads. Furthermore, high-quality reads were aligned to the improved reference assembly (Cajca.Asha_v2.0) (Garg et al., 2022) using BWA version 0.5.9 (Li et al., 2009) with the standard parameters. SNP calling was conducted via GATK v.3.7 (McKenna et al., 2010). Biallelic SNPs exhibiting less than 20% missing calls and a minor allelic frequency cut-off of 5% and 50 % heterozygosity were utilized for further analysis.

### 2.5 Linkage disequilibrium (LD) decay, GWAS analysis and candidate gene identification

LD was analyzed using TASSEL 5.0. The default parameters of PopLDdecay 3.4.2 were employed to calculate the decay of LD with physical distance. GWAS analysis was performed using 4,99,980 polymorphic SNPs and three seasons pooled phenotyping data recorded on PBR and component traits. In our study, we conducted GWAS analysis employing four models: MLM, CMLM, FarmCPU, and SUPER., utilizing R/GAPIT 4.3.1. The “Bonferroni correction” p-value threshold (<1.00004E-07) was implemented to remove false associations and only MTAs with a PVE > 0% were considered (**Supplementary Table 1)**. However, we found that significant MTAs were only identified using the FarmCPU and SUPER models, which provided the most reliable and statistically significant results, so we considered the MTAs from these two models for downstream analysis. The physical position of significant MTAs with associated traits were used to mine candidate genes in these regions on the pigeonpea reference genome assembly v2.0 (Garg et al., 2022). We only considered genes where significant MTAs were located (genic and non-genic region), based on variant annotation and the prediction of SNP effects using the open source SNPEff-4.3T program.

## 3. Results

### 3.1 Phenotypic variation, heritability and correlation for PBR and component traits

Phenotypic evaluation for PBR and component traits showed significant variation among the minicore accessions. A symmetric distribution was observed for most of traits (**Figure 1**). PBR score showed differences across seasons (3-9 in S1; 4-9 in S2; and 5-9 in S3). The average score recorded were 7.3, 6.5, and 7.3, respectively. Similarly, accessions revealed a wide range of variation for component traits; for DF (48-185 in S1; 50-180 in S2 and 70-180 in S3); DM (90-245 in S1, 95-230 in S2, 115-230 in S3); and GY (3-287 g plant^-1^, 8-463 g plant^-1^ and 6-481 g plant^-1^). Across seasons, all traits showed a larger phenotypic coefficient of variation (PCV) and genotypic coefficient of variation (GCV) (>□10%) respectively. The broad sense heritability (h^2^) averaged 54% for PBR, 92 % for DF, 96 % for DM and 55% for GY (**Table 1**). Following the replicated multi-season field evaluation results, 19 best lines were selected and screened using lab-based pod bioassay. The gain % compared to the resistant check (ICPL 332; score 6) and susceptible check (ICPL 87; score 9) was assessed. Among the best lines, four lines-ICP 10503, ICP 655, ICP 9691 and ICP 9655 (scoring between 4-5)-showed the least damage rating and low larval weight. Pearson’s correlation test was performed to determine the phenotypic correlation between PBR and component traits. A total of 6 possible correlations were observed, with 3 pairs (1 positive and 2 negative). Correlation discussion revealed a strong positive correlation between the DM and DF (r□=□0.97), whereas the GY and PBR had the highest negative correlation, (r□=□-0.55), which was significant at the 0.001 level. Other correlations were not statistically significant (p > 0.05). (**Figure 2**).

**Table 1.** Mean, range, and variability components in minicore accessions for PBR and component traits across three seasons.

| Traits | Seasons | Range | Grand Mean | CD (5%) | SE (±) | GCV | PCV | H<sup>2</sup> (%) |
| --- | --- | --- | --- | --- | --- | --- | --- | --- |
| **DF** | S1 | 48-185 | 118.5 | 14.8 | 5.3 | 17.9 | 19.5 | 80 |
|  | S2 | 50-180 | 123.7 | 7.5 | 2.7 | 18.3 | 18.5 | 97 |
|  | S3 | 70-180 | 125.6 | 3 | 1.1 | 13.6 | 13.6 | 99 |
| **DM** | S1 | 90-245 | 162.7 | 6.1 | 2.2 | 13.7 | 13.9 | 97 |
|  | S2 | 95-230 | 171.3 | 10.6 | 3.8 | 14 | 14.3 | 95 |
|  | S3 | 115-230 | 175.5 | 4 | 1.4 | 10.1 | 10.2 | 98 |
| **PBR** | S1 | 3-9 | 7.3 | 1.8 | 0.7 | 13.3 | 20.6 | 42 |
|  | S2 | 4-9 | 6.5 | 1.6 | 0.6 | 16.7 | 21 | 63 |
|  | S3 | 5-9 | 7.3 | 1.5 | 0.5 | 12.3 | 16 | 58 |
| **GY(gm)** | S1 | 3-287 | 124.1 | 144 | 34.9 | 57.4 | 76.9 | 56 |
|  | S2 | 8-463 | 168.5 | 154.9 | 55.4 | 43.7 | 63.8 | 47 |
|  | S3 | 6-481 | 216.7 | 129 | 46.2 | 40.4 | 50.4 | 64 |

**S1:** Rainy 2007; **S2:** Rainy 2008; **S3:** Rainy 2009; **DF:** Days to 50% flowering; **DM:** Days to maturity; **PBR:** Pod-borer complex resistance; **GY:** grain yield; **SE:** Standard Error; **CD:** Critical difference; **GCV:** Genotypic Coefficient of Variance; **PCV:** Phenotypic Coefficient of Variance; **H<sup>2</sup>:** Broad sense heritability

**Figure 1.**
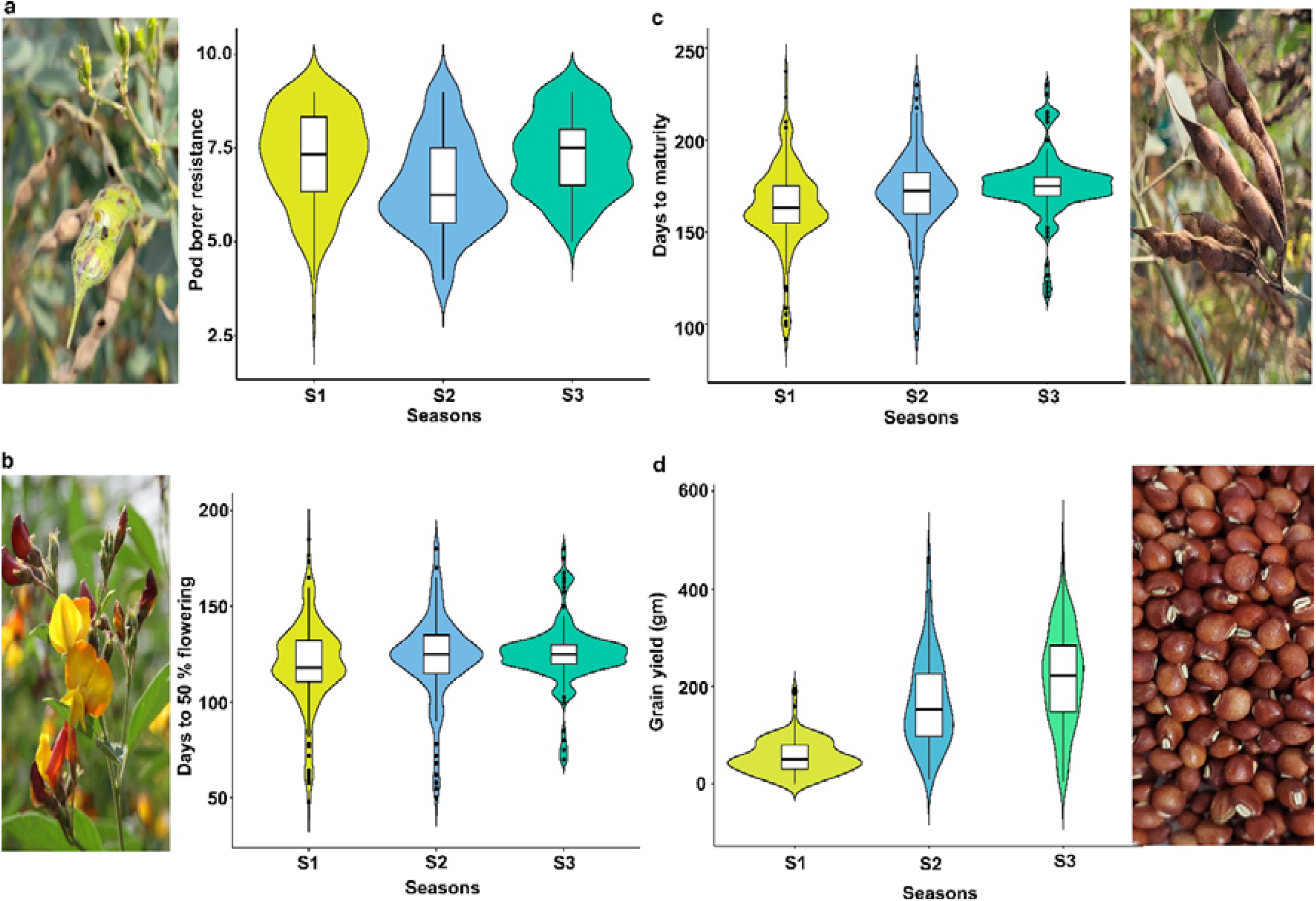
Phenotypic variation in minicore accessions for PBR and component traits: Violin plot showing variation for **(a)** Pod borer resistance (PBR); **(b)** Days to 50% flowering, **(c)** Days to maturity; **(d)** Grain yield (gm) traits consecutively evaluated for three seasons (**S1**: Rainy 2007; **S2**: Rainy 2008; **S3**: Rainy 2009)

**Figure 2.**
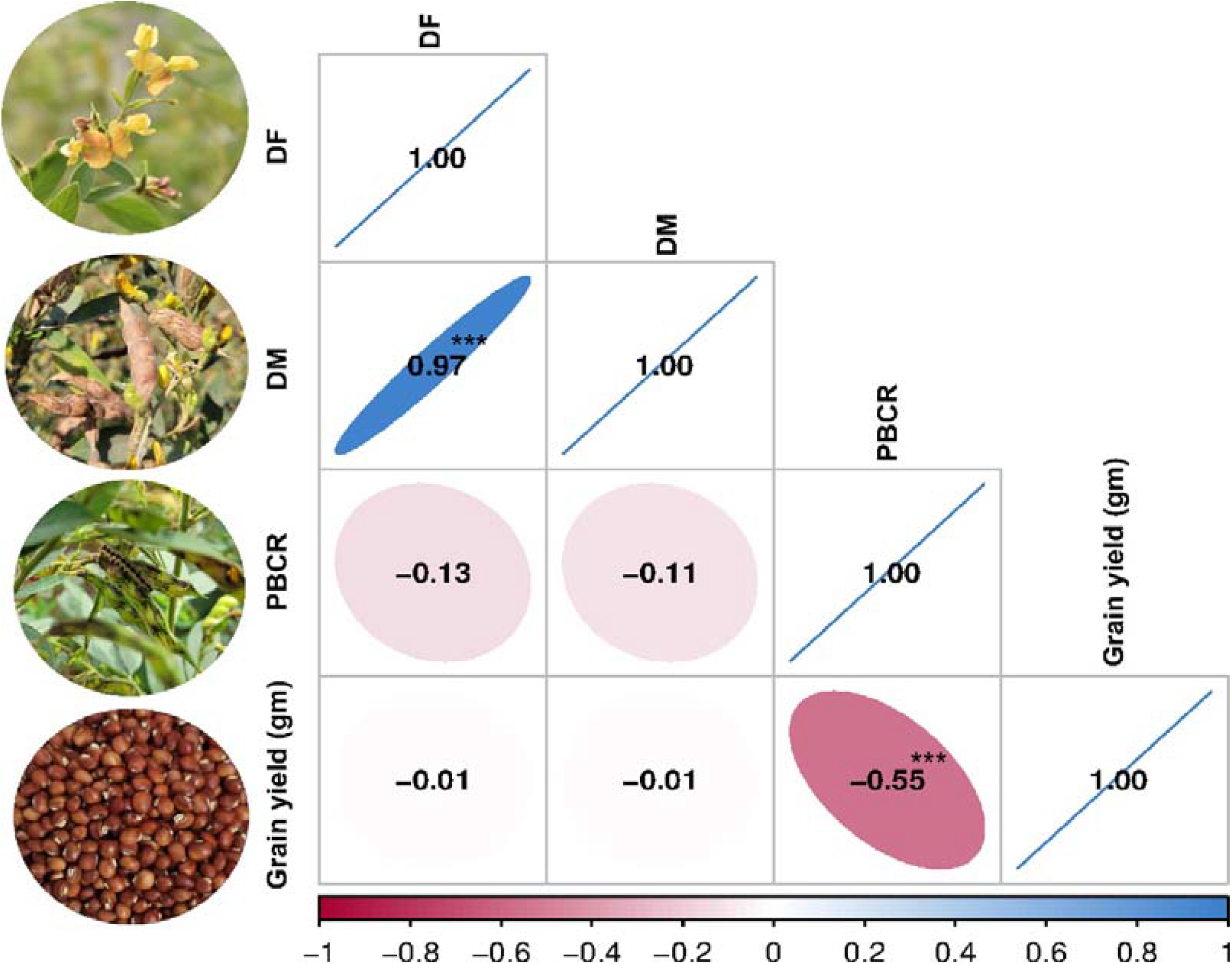
Pearson correlation matrix. Correlation between pod borer resistance (PBR) and component traits. (^***^ significant at 0.001 level)

### 3.2 LD decay and genome-wide association analysis

A total of 4,99,980 filtered SNP markers with a genotype call rate > 0.80 and a minor allele frequency (MAF) of >5% were utilized for downstream analysis. The overall LD decay across the 11 chromosomes was 39.5 kbp on average (**Figure 3**). For the GWAS analysis, high-quality phenotyping data (three seasons pooled phenotyping data) for PBR and component traits, were used along with 4,99,980 polymorphic SNPs. Two models (SUPER, FarmCPU) identified 14 significant MTAs, (8 PBR, 1 DF, 2 DM and 3 GY) respectively, for four traits, explaining 0.05%–28.1% phenotypic variation with a *p*-value range of 1.10E-13 to 9.66E-09 for pooled data (**Table 2**). Eight significant MTAs were identified for PBR on chromosomes CcLG02, CcLG04, CcLG05, CcLG07, and CcLG11, with PVE ranging from 0.05–5.57% (**Figure 4**). For DF, one MTA was detected (CcLG11_38698041) on the same chromosome (CcLG11) with PVE of 28.1%. On chromosome CcLG04, two MTAs (CcLG04_38227177 and CcLG04_7181399) for DM were detected, explaining 1.07–2.74% PVE. Three MTAs were identified for GY on chromosomes CcLG02, CcLG05, and CcLG07, accounting 0.49–1.72% phenotypic variance (Figure 5). Based on pooled phenotyping data, a representative set of minicore accessions, exhibiting variability for PBR and component traits, was selected to in silico validate the SNPs associated with the significant MTAs. Among the 14 detected MTAs, 5 showed polymorphism, including two for PBR (CcLG04_35844765 and CcLG07_10581882) (**Supplementary Figure 2**) and three for component traits (1 for DF: CcLG01_38698041; 1 for DM: CcLG04_7181399; and 1 for GY: CcLG04_25609089) in the minicore accessions (**Supplementary Figure 3-5**). These results indicate that identified SNPs could be used to develop allele-specific markers for MAS, helping develop pigeonpea cultivars with improved resistance to *H. armigera*.

**Table 2.** Significant MTAs detected for PBR and component traits using multi-locus models with predictive gene variants and their functions.

| Traits | Significant MTAs | Models | Chr | Position<br>(bp) | *p*-value | PVE<br>(%) | Gene<br>variants | Alleles | Gene ID | Functional Annotation |
| --- | --- | --- | --- | --- | --- | --- | --- | --- | --- | --- |
| PBR | CcLG02_16059144 | SUPER | CcLG02 | 16059144 | 8.56E-09 | 1.52 | DGV | G/A | Cc_3152 | *Probable carboxylesterase 15* |
| PBR | CcLG04_31940898 | SUPER | CcLG04 | 31940898 | 6.40E-09 | 1.30 | DGV | T/C | Cc_10409 | *Microtubule-associated protein 5* |
| PBR | CcLG04_35844765 | SUPER | CcLG04 | 35844765 | 9.55E-09 | 5.57 | UGV | G/A | Cc_10514 | *Phospholipase D delta* |
| PBR | CcLG05_29876072 | SUPER | CcLG05 | 29876072 | 2.21E-08 | 3.35 | 5'UTR | A/T | Cc_11847 | *Uncharacterized protein LOC100796483* |
| PBR | CcLG05_10183191 | SUPER | CcLG05 | 10183191 | 9.66E-09 | 0.31 | UGV | C/A | Cc_11087 | *L-type lectin-domain containing receptor kinase IX.1* |
| PBR | CcLG05_9948484 | SUPER | CcLG05 | 9948484 | 2.13E-08 | 0.15 | UGV | C/A | Cc_11074 | *Hypothetical protein GLYMA_14G106500* |
| PBR | CcLG07_10581882 | SUPER | CcLG07 | 10581882 | 1.95E-11 | 0.05 | UGV | G/A | Cc_15562 | *Omega-hydroxypalmitate O-feruloyl transferase* |
| PBR | CcLG11_49001007 | SUPER | CcLG11 | 49001007 | 7.86E-08 | 4.48 | SV | T/C | Cc_23491 | *FAR1-RELATED SEQUENCE 4* |
| DF | CcLG11_38698041 | SUPER | CcLG011 | 38698041 | 5.96E-08 | 28.11 | UGV | C/T | Cc_23683 | *Hypothetical protein LR48_Vigan05g175700* |
| DM | CcLG04_38227177 | FarmCPU | CcLG04 | 38227177 | 5.53E-08 | 1.07 | UGV | G/T | Cc_10599 | *RING-H2 finger protein ATL7* |
| DM | CcLG04_7181399 | FarmCPU | CcLG04 | 7181399 | 1.74E-11 | 2.74 | UGV | G/A | Cc_9357 | *Leucine-rich repeat receptor-like protein kinase* |
| GY | CcLG02_25609089 | FarmCPU | CcLG02 | 25609089 | 1.10E-13 | 0.49 | UGV | C/T | Cc_3417 | *Glutaredoxin-C5, chloroplastic* |
| GY | CcLG05_16338260 | FarmCPU | CcLG05 | 16338260 | 1.75E-12 | 1.72 | DGV | C/G | Cc_11279 | *Secretory carrier-associated membrane protein* |
| GY | CcLG07_9297796 | FarmCPU | CcLG07 | 9297796 | 1.22E-11 | 0.97 | UGV | C/T | Cc_15498 | *Hypothetical protein GLYMA_16G039300* |

**MTAs:** Marker trait associations; **Chr:** Chromosome; **PBR:** Pod-borer complex resistance; **DF:** Days to flowering; **DM:** Days to maturity; **GY:** Grain yield; **UGV:** Upstream gene variant; **DGV:** Downstream gene variant; **5'UTR:** 5 prime UTR variant; **SV:** Synonymous variant; **MGV:** Missense gene variant; **PVE:** Phenotypic variation explained

**Figure 3.**
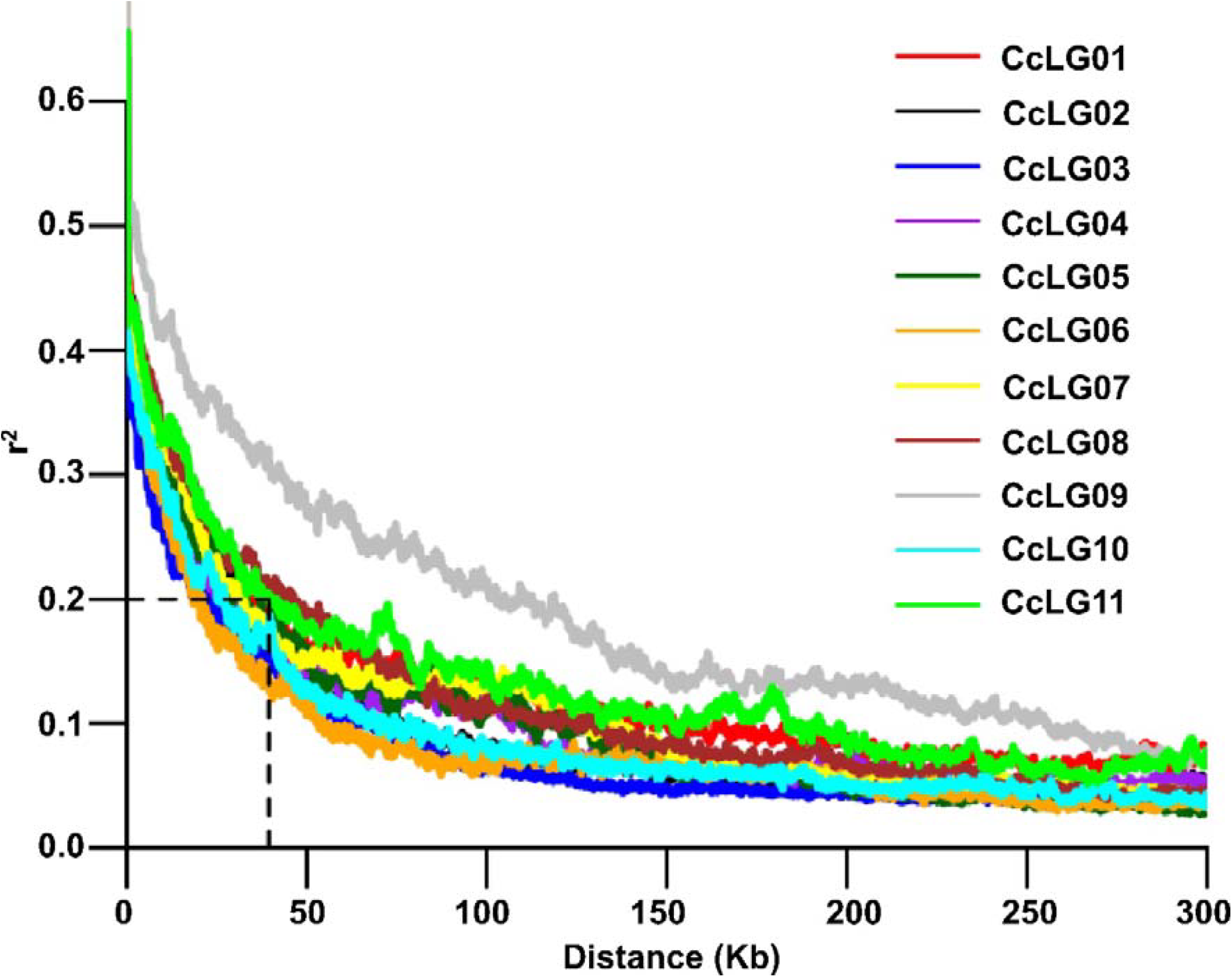
Estimated linkage disequilibrium decay (LD-decay). LD decay for each chromosome, with r^2^ ~ 0.2 at 39.5 kb

**Figure 4.**
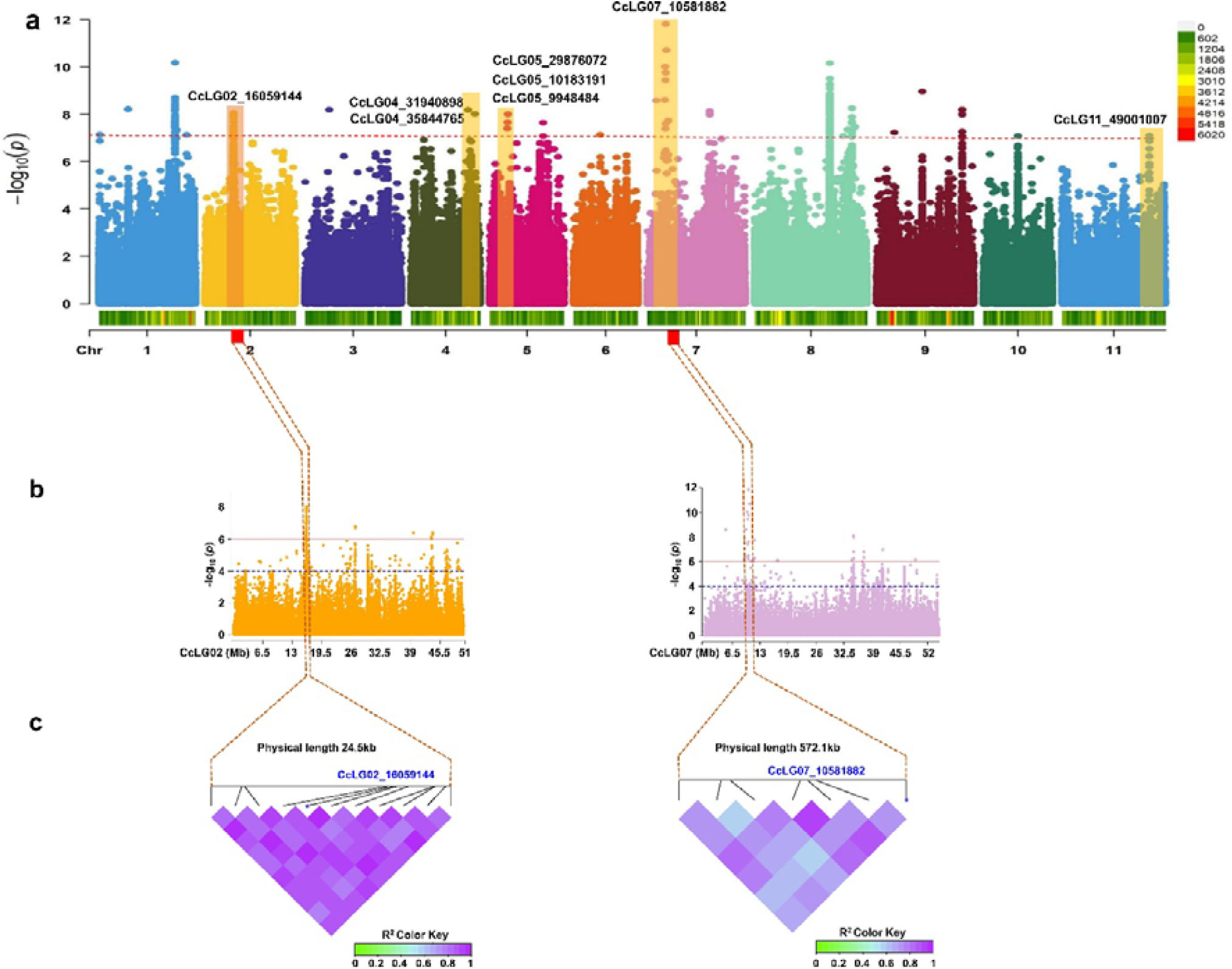
Genome□wide association study (GWAS) for pod borer resistance (PBR) trait: **(a)** Manhattan plot illustrating significant MTA for PBR trait. Only highly statistically significant MTAs at peak were considered; **(b)** Association for the significant SNPs on chromosomes CcLG02 and CcLG07. The interval of association was determined to lie between 24.5 kb downstream of the significant MTA (CcLG02_16059144) on CcLG02 and 572.1 kb downstream of the significant MTAs (CcLG07_10581882) on CcLG07; **(c)** Linkage disequilibrium heatmaps for the association region for PBR on chromosome CcLG02 and CcLG07. Bonferroni correction threshold of *p-*value (<1.00004E-07) was implemented to detect significant associations.

**Figure 5.**
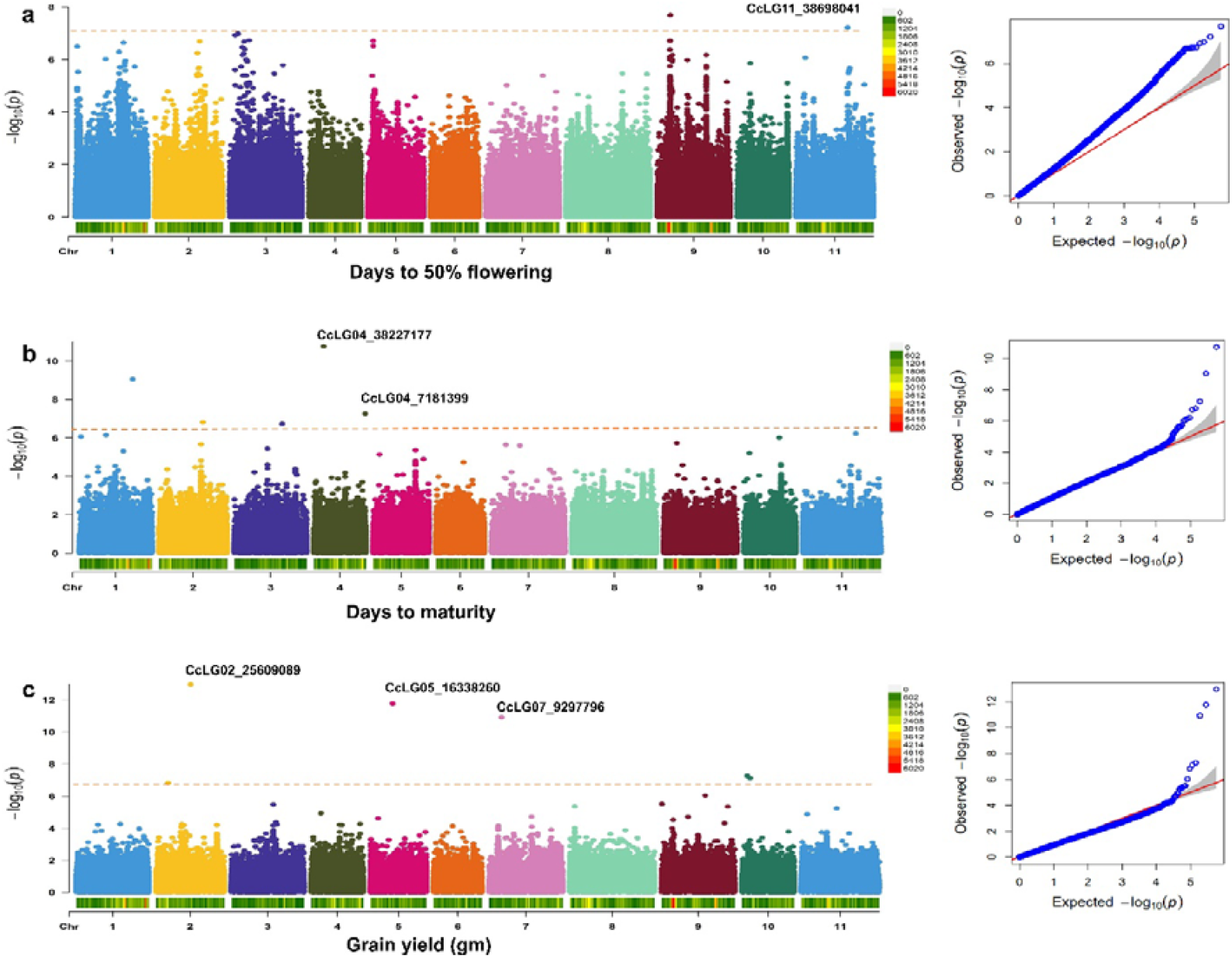
Genome□wide association study (GWAS) for component traits: Manhattan and quantile-quantile (QQ) plots for **(a)** Days to 50% flowering; **(b)** Days to maturity; and **(c)** Grain yield. Bonferroni correction threshold of *p-*value (<1.00004E-07) was implemented to detect significant associations.

### 3.3 Putative genes associated with MTAs

We examined the putative genes linked with the 14 significant MTAs identified for PBR and component traits by analyzing their location (genic and non-genic), effects, and functions (**Table 2**). Among these, 12 were detected in the intergenic regions, while one each was found in the exonic, 5′ UTR, and synonymous variant regions. Notably, the MTA CcLG05_29876072, located in the 5′ UTR of gene (Cc_11847) on CcLG05 chromosome, was associated with PBR. Additionally, the MTA-CcLG11_49001007, located in the synonymous variant region of gene (Cc_23491) on CcLG11 chromosome, was also associated with PBR. Furthermore, the remaining significant MTAs associated with PBR, DF, DM and GY were found in intergenic regions, respectively.

## 4 Discussion

Pigeonpea is a major grain legume that draws worldwide interest for its important contribution to nutritional and food security. However, *H. armigera* results in substantial losses in yield, posing a severe obstacle in pigeonpea cultivation (Sharma et al. 2022). Although significant breeding efforts have been made, the development of resistant varieties remains difficult due to the complex inheritance of resistance traits and the lack of genetic variation in cultivated germplasm (Volp et al. 2022, Karrem et al., 2025). Whereas transgenic approaches have potential, legal issues and public acceptability concerns in India (Rakesh and Ghosh 2024). Moreover, conventional pest management practices often lack sustainable solutions, due to resistance to pesticides and environmental issues. The investigation of pigeonpea germplasm in the primary gene pool and crop wild relatives is an effective option, since several *Cajanus* species exhibit higher resistance to *H. armigera* (Kumari et al. 2010; Sharma et al. 2022; Singh et al., 2022). However, most cultivated genotypes showed low to moderate levels of resistance to *H. armigera*, evidenced by the screening of nearly fourteen thousand pigeonpea accessions (Reed and Lateef 1990). Several investigations reported, few accessions of the wild progenitor of pigeon pea have exhibited high levels of resistance to *H. armigera* (Green et al., 2006; Sharma et al., 2009; Sharma et al., 2022). It is important to understand the trait is substantially influenced by genetic and environmental factors; therefore, relying solely on phenotypic screening for selection is insufficient. Furthermore, understanding the genetic basis of resistance to *H. armigera* can provide opportunities for developing resistance varieties. Our investigation on minicore accessions reported broad range of variation in the PBR and component traits. The broad sense heritability (h^2^) of PBR, DF, DM, and GY were 54%, 92%, 96%, and 55%, respectively, indicating a significant portion of the variation is attributable to distinct genotypes. The correlation analysis revealed a strong positive correlation between DM and DF, indicating that days to flowering could act as an index for maturity classification in pigeonpea. Previous investigations showed the similar findings for a correlation between DM and DF (Singh et al., 1995). Compared to other traits, GY and PBR had the strongest negative association (r = −0.55), which shows highly influential nature of trait. This indicates higher PBR score tends to associated with lower GY, or vice versa. Furthermore, environmental factors such as excessive rainfall during sowing in S1, delayed planting, and variations in day and night temperatures throughout the reproductive stages likely contributed to the lower yield in S1, despite similar pod borer scores in other seasons. Previous studies have also reported that delayed sowing reduces yield in pigeonpea (Arunkumar et al., 2018).

MAS is an intriguing approach for faster up the development of insect pest’s resistant varieties. It facilitates the development of multi-trait resistant varieties by pyramiding different resistance genes to target insects, that is not possible with traditional breeding due to alike expression of phenotype (Sharma & Crouch, 2004). Utilizing WGRS data along with precise phenotypic variability could help identify accessions with rare variants that may be potentially linked with important traits, such as resistance to *H. armigera*. In GWAS, determining the pattern of LD is important, since it influences the resolution and magnitude of the association analysis. Our analysis showed an average LD decay at 39.5 kbp. The rapid LD decay indicates a minimal extent of long-range LD among the minicore accessions. Previous study reported, genome-wide LD decay at 118 kb (Megha et al., 2024). GWAS minimizes the two primary constraints of traditional linkage mapping, such as limited allelic diversity and insufficient genetic resolution (Huang and Han, 2014). Due to its high resolution, and low cost in sequencing/genotyping, GWAS analysis has successfully dissect important traits in pigeonpea, including flowering related traits (Kumar et al., 2022); antioxidant activity (Megha et al., 2024). This aspect can facilitate fine-scale association mapping for finding genetic variations linked to particular phenotypic traits. The main concern for GWAS is to minimize false positives, mostly due to population structure and familial relatedness (Kaler et al., 2020). Although single-locus models overcome this issue by including the two confounding factors as covariates, over fitting in a model usually leads in false-negatives, which could eliminate valuable loci (Price et al., 2006). In this context, multi-locus models provide an alternative way for reducing false negatives (Zhang et al., 2019). Multi-locus GWAS models, such as settlement of MLM under progressively exclusive relationship (SUPER) and fixed and random circulating probability unification model (FarmCPU) methods improve statistical power although minimizing false positives. The SUPER model offers greater computational power and requires less computing than earlier models. However, it extracts a small number of SNPs termed pseudo quantitative trait nucleotide (QTN) to determine kinship (Wang et al., 2014). Moreover, the “FarmCPU” is a novel multi-locus model that is computationally powerful and efficiently controls false negatives and false positives. Two multi-locus methods (SUPER and FarmCPU) were in the current investigation to identified significant MTAs for PBR and component traits. GWAS analysis identified 14 significant MTAs linked to 4 traits, including 8 for PBR, 3 for GY, 2 for DM and 1 for DF. For DF trait, we detected one MTAs on chromosome CcLG11, accounting highest phenotypic variation 28.1%. Most of the identified MTAs exhibited smaller phenotypic variation % and lower P-values. This finding suggests that these traits are controlled by multiple genes with minor effects, reflecting complex genetic architecture and are also influenced by environmental factors. The statistical power of association mapping could be substantially improved by increasing the population size (Liu et al., 2021). The MTAs detected for PBR and component traits in our study were not reported previously and seems to indicate novel genetic loci in pigeonpea. Thus, the SNPs associated with MTAs offers the possibility of additional validation in diverse collections and might be utilized for early generation selection in breeding programs.

A total of 14 significant MTAs for four traits were detected and linked with putative genes. One MTA for PBR was found on chromosome 2 (CcLG02_16059144) linked to the *Cc_3152* gene encoding a *probable carboxylesterase 15* enzyme that catalyzes the conversion of carboxylic esters and water into alcohol and carboxylate. In plants, it is involved in defense, development, and secondary metabolism (Palayam et al., 2024). In tobacco, this gene (NbCXE) involved in host defense responses against TMV infection (Guo et al., 2020). Similarly, another MTA (CcLG04_31940898) was identified for PBR encoding *microtubule-associated protein 5* which play key role in cell division, cell proliferation, and cell morphology. In Arabidopsis, the microtubule-binding protein (*TGNap1*) facilitates the secretion of antimicrobial proteins, important for defense against phytopathogens (Bhandari et al., 2023). The MTA detected for PBR (CcLG11_49001007) in the exonic region of the gene *Cc_23491*, which encodes *FAR1-RELATED SEQUENCE*, is a light signaling factor pair with *FAR-RED ELONGATED HYPOCOTYL* 3 to regulate plant immunity by integrating chlorophyll biosynthesis with the salicylic acid (SA) signaling pathway in Arabidopsis (Wang et al., 2015). For PBR, three more MTAs were detected and associated with *Cc_10514, Cc_11087*, and *Cc_15562*. The gene *Cc_10514* encodes *phospholipase D delta*, a protein that involved in basal defense and non-host resistance to powdery mildew fungi in Arabidopsis (Pinosa et al., 2013). *Cc_11087* encodes an L-type lectin domain-containing receptor kinase IX, involved in self/non-self-surveillance and plant resistance. The homologues of these receptors in *Nicotiana benthamiana* and *Solanum lycopersicum* have same role in defense against Phytophthora (Wang et al., 2015) and MTA identified for PBR on chromosome 7 (CcLG07_10581882) associated with *Cc_15562* gene encoding *Omega-hydroxypalmitate O-feruloyl transferase*, role in suberin biosynthesis. Suberin is synthesized in plant wound tissues to prevent pathogen infection (Molina et al., 2009). For PBR, two MTAs (CcLG05_29876072 and CcLG05_9948484) are detected in exonic (*Cc_11087* gene) and intergenic (*Cc_11074* gene) regions, respectively. These are predicted to encode an *uncharacterized protein* and a *hypothetical protein GLYMA_1*.

The MTA (CcLG04_38227177) identified for DM lie with the *Cc_10599* gene, encodes for *RING-H2 finger protein* which is important for seed development in Arabidopsis (Xu et al., 2003). The MTA (CcLG04_7181399), identified for DM, is associated with the *Ca_00148* gene, which encodes a *leucine-rich repeat receptor-like protein kinase*. This protein is a important membrane-bound regulator of abscisic acid (ABA) early signaling in Arabidopsis, and ABA involved in seed maturation (Osakabe et al., 2005). For GY, *CcLG05_25609089* MTA was associated with *Cc_3417* gene. The *Cc_3417* encodes a *glutaredoxin-C5 chloroplastic protein*. Overexpression of a *CPYC-type glutaredoxin* was shown to increase grain weight in rice (Liu et al., 2019). The another MTA (CcLG05_16338260), is present in the intergenic region of Cc_11279 gene, encodes for the *Secretory carrier associated membrane protein*. Karnik et al. (2013) have demonstrated that SCAMPs are involved in the secretion of defence proteins, including protease inhibitors and toxins, in Arabidopsis thaliana. These proteins have been shown to inhibit insect feed or growth. However, further validation of the identified MTAs is required across varying genetic backgrounds. This provides deeper insights into the genetic control of resistance mechanism, along with the potential to develop effective markers (Thakur et al., 2025). Additionally, gene editing innovations, offer promising tools for validating and modifying the candidate genes identified by GWAS. It enables the precise knock-in or knockout of specific genes, providing clear evidence of their role in resistance. The integration of detected genes and SNPs associated with MTAs through molecular breeding or genetic modification could provide an effective approach for developing *H. armigera*-resistant cultivars.

## Conclusion

Pod borer, *H. armigera*, is one of most damaging pest in pigeonpea production. Various methods have been employed for controlling this pest, exhibit limited success. Phenotypic data on PBR and component traits, along with genotypic data from the WGRS, were used to identify 14 significant MTAs. These significant MTAs, explaining 0.05%–28.1% phenotypic variation with *p* value range of 1.10E-13 to 9.66E-09. MTA for DF (CcLG11_38698041) on chromosome (CcLG11) explaining highest PVE of 28.1%. Furthermore, we identified important genes that encode *probable carboxylesterase 15* (*Cc_3152*), *microtubule-associated protein 5* (*Cc_10409*), and *FAR1-RELATED SEQUENCE* (*Cc_23491*) have been associated with plant defence responses and regulation of plant immunity. These putative genes can be helpful for the identification of molecular targets, providing insight into the biological pathways that underlie the traits of interest, and facilitating understanding of the genetic basis of complex traits. The importance of these genomic regions for future studies will help to understand the *H. armigera* resistance mechanism along with finding functional markers. Notably, further lab-based pod bioassay screening identified four minicore accessions—ICP 10503, ICP 655, ICP 9691, and ICP 9655—which showed moderate resistance. The resistant genotypes, significant MTAs, and putative genes identified in this investigation have the potential to be utilised in the development of pod borer–resistant pigeonpea cultivars.

## Author contributions

Conceptualization – MKP and HCS; resources and project administration – MKP, HCS and JJ; investigation and methodology AM, RSM & JJ; data curation and formal analysis AM, SPM, VS; Writing original draft AM, VS; reviewed and edited the draft – VS, SSG (Shailendra Singh Gaurav), SSG (Sunil S. Gangurde), ND, SR, RR, JJ, HCS and MKP. All the authors contributed to the article and approved the submitted version.

## Acknowledgments

AM acknowledges Chaudhary Charan Singh University (CCSU), Meerut, for collaborating with ICRISAT

## Data availability

All the data generated in the present study is provided in the Supplementary Information and sequencing data deposited as Bioproject PRJNA575817, PRJNA383013, PRJNA57990 in NCBI

## Declarations

### Ethics approval and consent to participate

Not applicable

### Consent for publication

All the authors have approved the submitted version of this manuscript for publication.

