## Supplementary Figures for "Dissecting genomic regions and candidate genes for pod borer resistance and component traits in pigeonpea minicore collection"

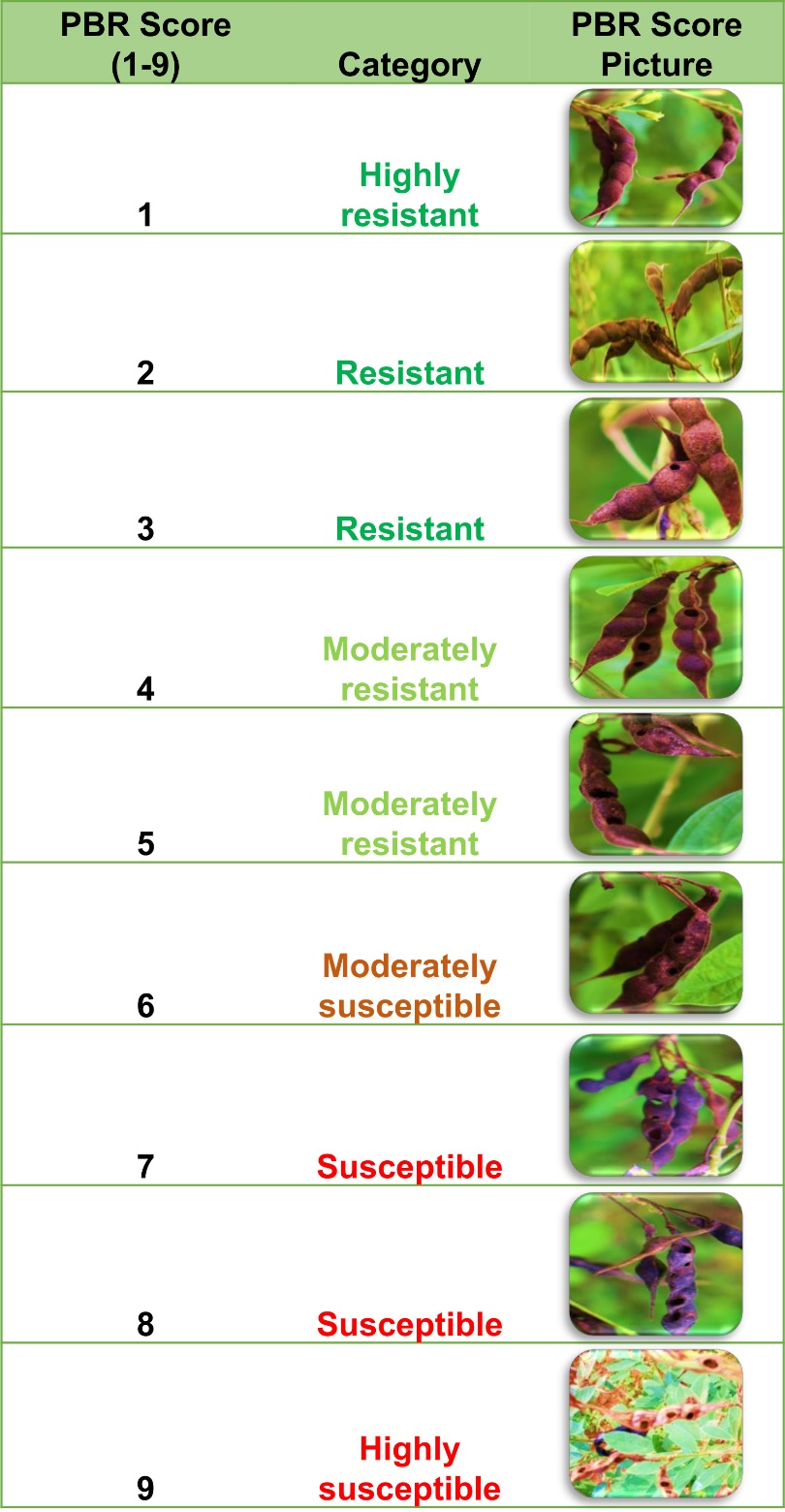

**Supplementary Figure 1. Disease Rating Scale**

**Pod borer complex resistance (PBCR) supplementary figures**

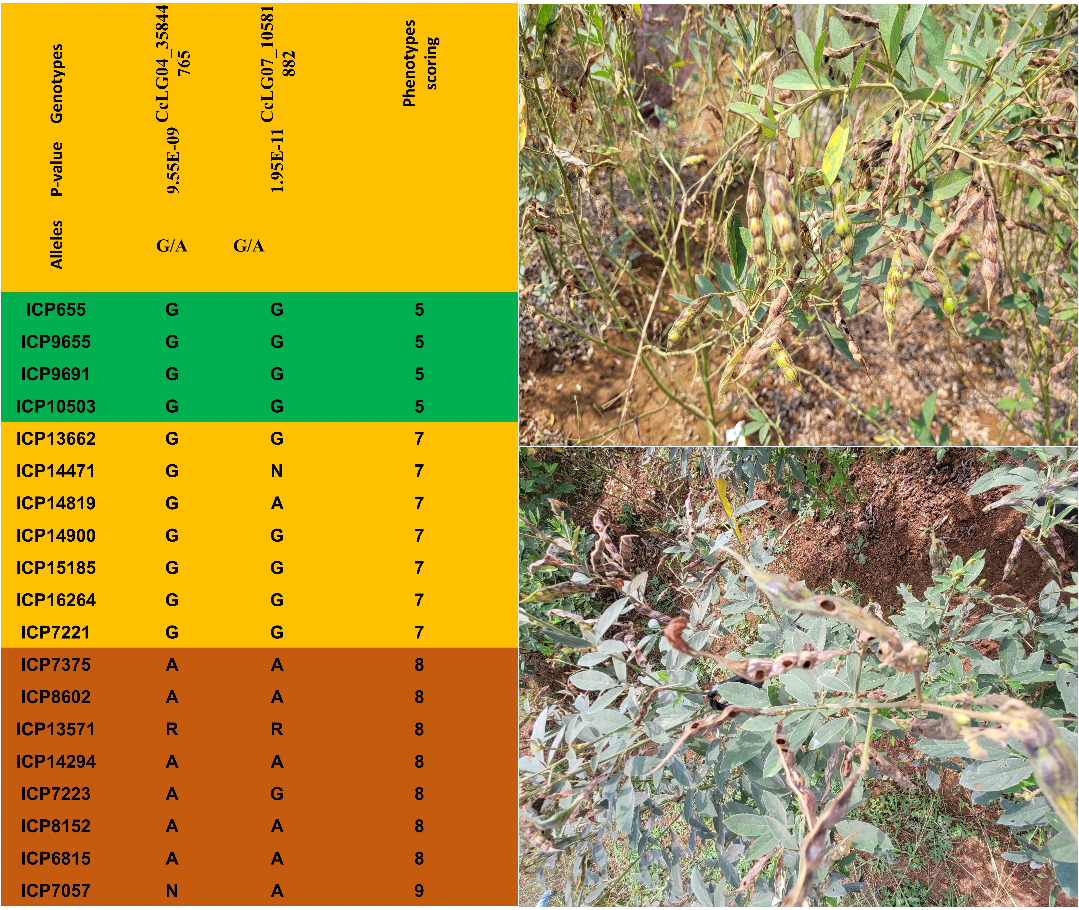

**Supplementary Figure 2.** GWAS results and allele effects of significant marker‒trait associations identified for PBCR. A) SNP marker loci and p values from GWASs for complex pod borer resistance. represented by the selected accessions corresponds to the allelic variation in the identified genomic regions. Phenotype of moderate resistance; susceptible phenotype.

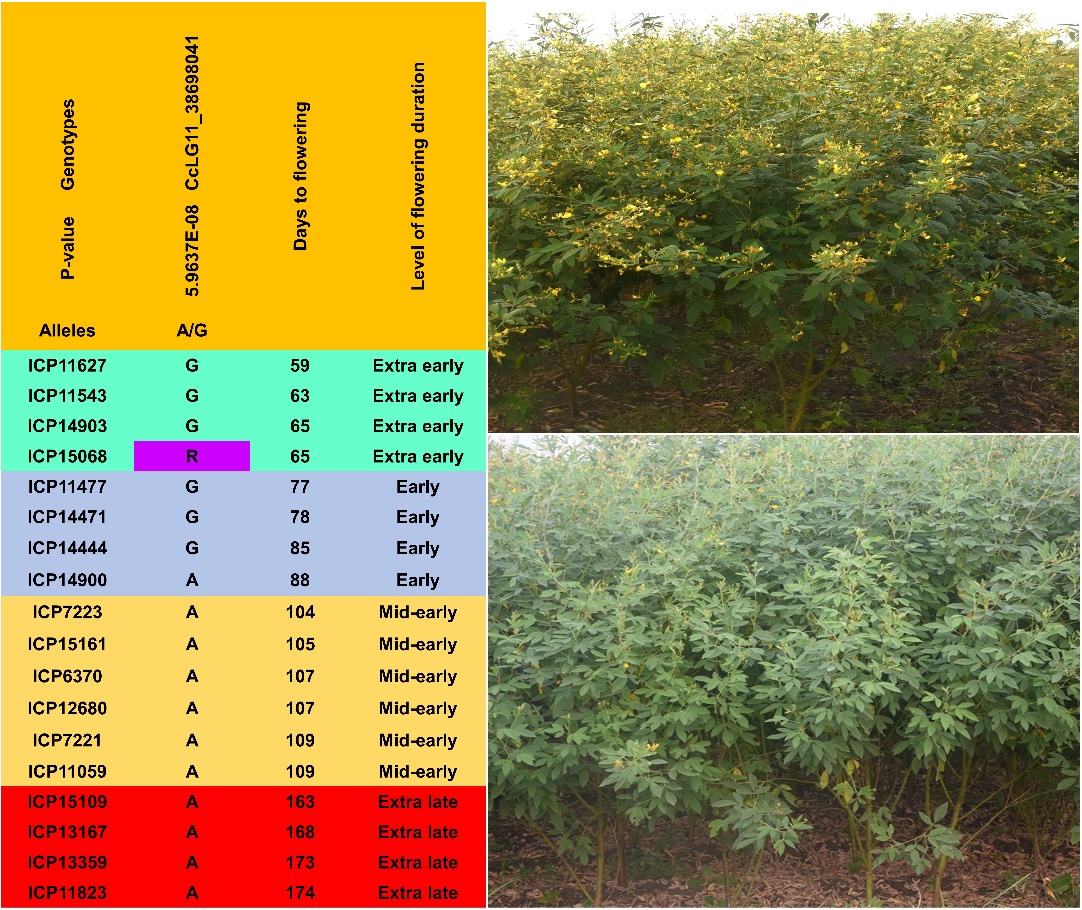

**Supplementary Figure 3.** GWAS results and allele effects of significant marker‒trait associations identified for days to flowering. represented by the selected accessions corresponds to the allelic variation in the identified genomic regions. Phenotype of early flowering; phenotype of late flowering.

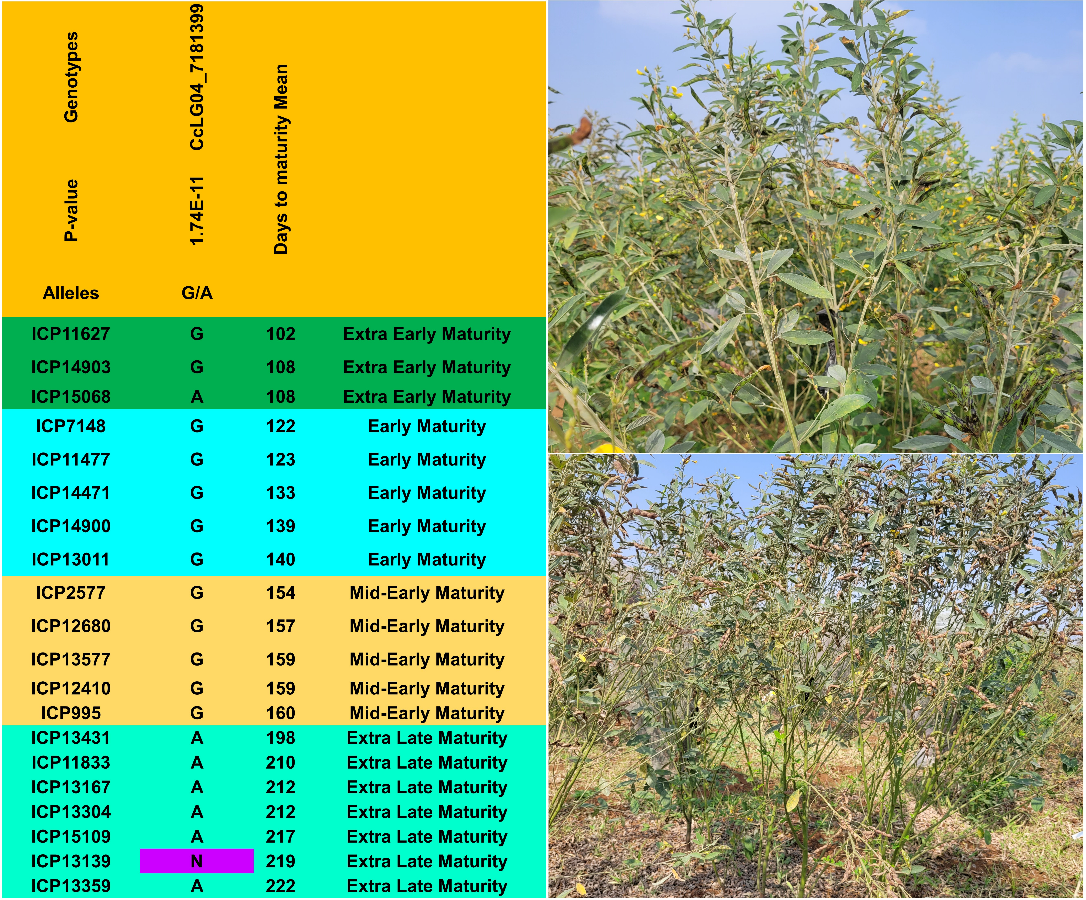

**Supplementary Figure 4.** Candidate single-nucleotide polymorphism (SNP) marker loci and p values from GWAS for days to maturity. The contrasting phenotype, i.e., early and late maturity, represented by the selected accessions corresponds to allelic variation in the targeted genomic regions.

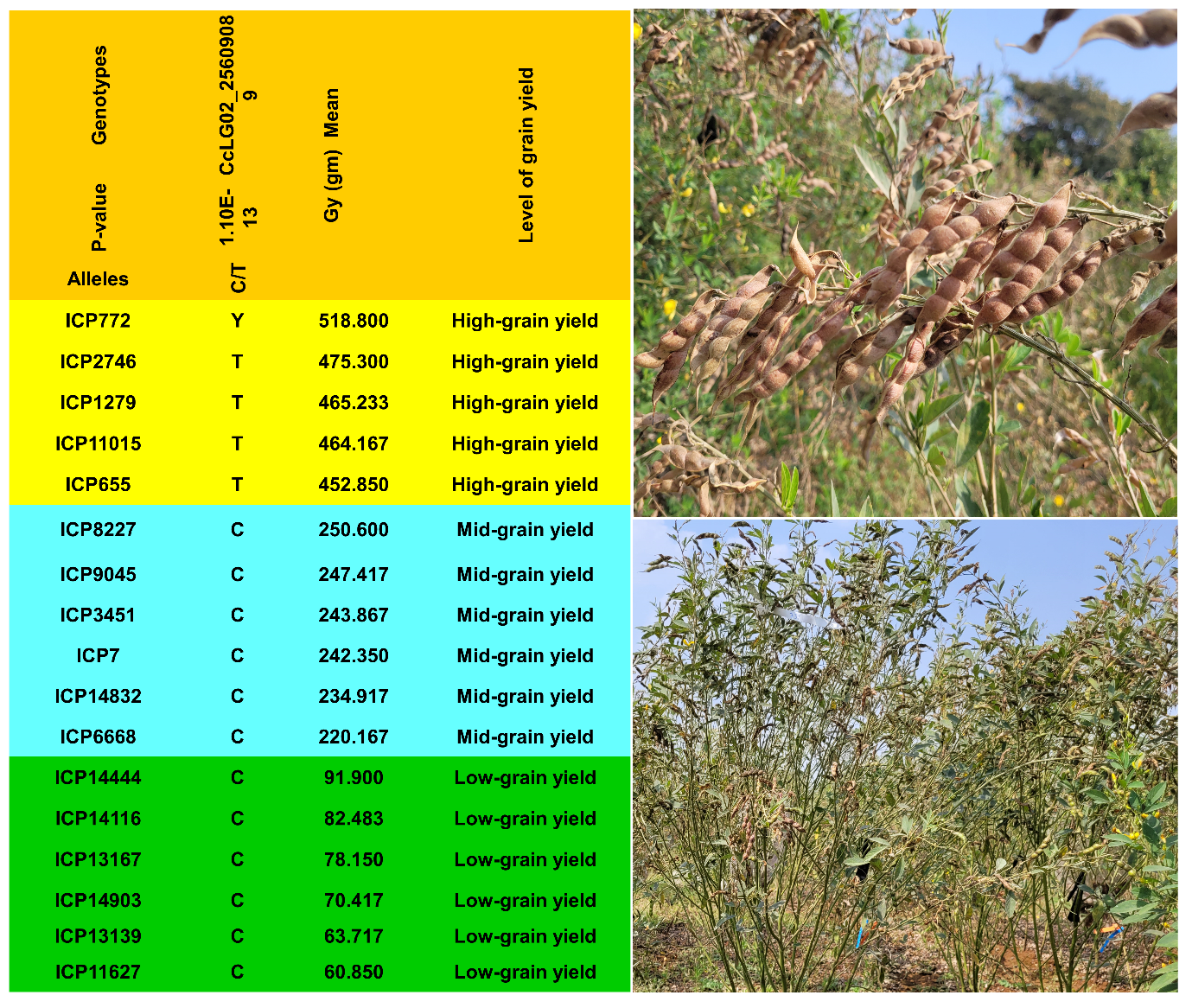

**Supplementary Figure 5.** Candidate single-nucleotide polymorphism (SNP) marker loci and p values from GWASs for grain yield. The contrasting phenotypes, i.e., high grain yield and low grain yield, represented by the selected accessions correspond to allelic variation in the targeted genomic regions.
